# Elusive scent of fear: no evidence of olfactory reversal conditioning in humans

**DOI:** 10.1101/2025.05.08.652858

**Authors:** N.S. Menger, B. Kotchoubey, Y.G. Pavlov

## Abstract

Reversal learning offers a window into how associations are acquired, updated, and overwritten. Because olfactory inputs bypass much of the thalamus and are tightly linked to emotion, we examined whether humans can flexibly form and subsequently reverse aversive associations to smells. Thirty healthy adults underwent an olfactory reversal conditioning protocol in which one neutral odor (CS+) was followed by a 90 dB aversive sound (US) and a second odor (CS-) was not. After five blocks the contingencies were reversed. Throughout 300 trials we collected ratings of pleasantness and intensity together with autonomic physiological indices (skin conductance, ECG, photoplethysmography, respiration), facial EMG, and 64-channel EEG. Contrary to expectations, pleasantness, intensity, and all autonomic or facial muscle measures failed to differentiate CS+ and CS-either before or after reversal (all *p* > .01). Event-related potentials, alpha suppression, heart rate and pulse wave responses likewise showed no CS specificity. Only multivariate classifiers: trained (i) on the time-domain EEG signal and (ii) on alpha-band activity, separately, distinguished CS+ from CS-at above-chance levels in late post-stimulus intervals. These neural signatures did not translate into overt physiological or behavioural differences. The pattern suggests that, at least with neutral odors and an auditory US, olfactory fear learning is subtle, spatially variable across the cortex, and easily masked in group-level averages. Our findings highlight both the promise of multivariate EEG for detecting fragile olfactory associations and the challenge of eliciting robust conditioned responses with cross-modal (odor-sound) pairings.

## Introduction

Classical conditioning is a well-established paradigm to examine basic forms of learning in humans. In fear conditioning, a neutral conditioned stimulus (CS) is followed by an unconditioned stimulus (US) that evokes a fear response. Through associative processes, the emotional properties of the US transfer to the CS. Once the association is formed, presenting the CS alone elicits a fear response, which is referred to as the conditioned response (CR). To control for habituation and physical stimulus characteristics, most paradigms employ two CS with similar features: a CS+ that is paired with a US, and a CS-which is not.

In addition to understanding how associations between CS and US are learned, it is crucial to investigate how they can be unlearned, which is an important area for understanding therapeutic interventions in fear-related disorders. Typically, this process is studied via an extinction phase following conditioning, during which the CS is presented repeatedly without the US, leading to a decrease in the CR. Another paradigm that can reveal the mechanisms of unlearning is reversal conditioning. In this approach, the roles of the stimuli are reversed after a set period: the original CS+ is no longer paired with the US, while the original CS-becomes paired. Successful performance therefore demands both suppression of the outdated memory and rapid formation of a new one - an index of cognitive flexibility and a closer analogue to therapeutic re-appraisal (Schiller et al., 2008).

The olfactory system presents a unique opportunity to probe such flexibility. Unlike vision or audition, olfactory afferents reach limbic and orbitofrontal regions with minimal thalamic relay, establishing an intrinsic connection between smell and affect. Many odors carry innate hedonic value, yet it remains unclear whether a neutral odor can acquire and, crucially, reverse a learned fear association as readily as visual or auditory cues. Empirical findings for successful fear acquisition learning are mixed, and for reversal are not present in the literature.

Although self-reported evaluations of a CS’s valence often remain unchanged after conditioning (Åhs et al., 2013; Cavazzana et al., 2018; Marinkovic et al., 1989), peripheral physiology frequently reveals conditioned responses. Most consistently, skin conductance rises to the CS+ (Gaby & Dalton, 2019; Leer et al., 2011; Marinkovic et al., 1989; Parma et al., 2015; Rosenthal et al., 2022). Respiratory findings are more variable: some studies report increased tidal volume to the CS+ (Devriese et al., 2006), others a decrease (Marinkovic et al., 1989), no change (Van den Bergh et al., 1997), or a transient effect that fades after a few trials (Leer et al., 2011). Respiration rate can likewise rise to the CS+ (Van den Bergh et al., 1997), although this difference is not always significant (Devriese et al., 2006). Heart-rate measures also provide partial evidence of conditioning (van den Bosch et al., 2015). Facial-muscle activity can differentiate odor valence outside a conditioning context (Armstrong et al., 2007; Kato & Yagi, 1994) and may serve a similar role here. Neural indices add information beyond autonomic arousal: Kastner et al. (2016) found no early difference in the late positive potential (LPP) between CS+ and CS-when odors acted as cues, but a significant divergence emerged later when the same odors served as contextual stimuli. Electro-olfactogram recordings show a conditioning-related shift in P1 latency, even without amplitude changes (Cavazzana et al., 2018). Finally, although not yet applied to olfactory paradigms, time-frequency EEG measures, such as CS-dependent changes in alpha suppression (Bacigalupo & Luck, 2022), might also prove sensitive to conditioned responses.

Physiological indices sometimes reveal clear differences between CS+ and CS-, whereas explicit ratings of pleasantness or intensity do not (Åhs et al., 2013; Cavazzana et al., 2018; Kastner et al., 2016; Marinkovic et al., 1989). To account for this mismatch in measurement sensitivity, we examined the neurophysiology of olfactory conditioning with an extensive battery of methods: throughout 300 trials we recorded ratings of pleasantness and intensity, skin conductance, electrocardiography, photoplethysmography, respiration, levator and corrugator electromyography and 64-channel EEG. We adopted a reversal paradigm to capture both learning and unlearning: an initially neutral odor (CS+) was paired with a loud aversive sound (US) while a second neutral odor served as CS-. Midway through the session the contingencies were switched, forcing participants to extinguish the original association and acquire a new one to the other odor.

The study pursued three specific aims: (1) to determine whether a neutral odor can form and subsequently reverse an aversive association when the US is auditory, (2) to compare the sensitivity of subjective ratings, autonomic measures, facial EMG and univariate EEG indices in tracking these associative changes and (3) to assess whether multivariate decoding of time-domain ERPs and alpha activity can detect conditioned differences that conventional analyses overlook.

## Methods

### Participants

In total, 48 participants were recruited via an email to all members of the University of Tübingen. Exclusion criteria were disorders of the nervous system, anosmia, pregnancy or breastfeeding, psychological disorders, and hearing disorders. Eight participants were excluded, two of whom due to technical issues, three due to exclusion criteria that were only identified when participants were already in the lab, and three aborted their participation. As it has been shown before that only aware participants demonstrate conditioning (Marinkovic et al., 1989), ten participants who were unable to report the CS-US contingencies were excluded from analyses, yielding a final sample of 30. Their mean age was 27.2 years (*SD* = 8.8, range 19 to 51), 66.7 % were female, and all were right-handed.

For specific physiological measures we discarded data with poor signal quality: two additional participants for heart rate, 13 for pulse wave amplitude, one for respiration rate, and two for EEG.

Participants were paid or received course credit for their participation. All participants gave informed consent before the experiment. The study followed the revised Declaration of Helsinki, and was approved by the ethics committee of the medical faculty at the Eberhard Karls University of Tübingen.

On the test day all participants rated their sense of smell as normal, with an average score of 71.93 (*SD* = 18.11) on a 0 to 100 scale where 0 represents “no smell” and 100 “excellent”.

### Stimuli

All odorants were diluted in mineral oil, leading to 20 ml liquids that were filled into the 100 ml jars in the olfactometer. The ratios of odorant and mineral oil were based on other olfactory studies (Amsellem et al., 2018; Höchenberger et al., 2015; Keller et al., 2012), and a pilot study: Orange oil (15 % v/v; CAS 8008-57-9), Lemon oil (5 % v/v; CAS 8008-56-8), Butyric acid (2 % v/v; CAS 107-92-6), Heptyl acetate (1 % v/v; CAS 112-06-1), Nonanal (5 % v/v; CAS 124-19-6), Linalool (1 % v/v; CAS 78-70-6), Mineral oil (100 % v/v; CAS 8042-47-5). The neutral odors that were open to selection as CS were heptyl acetate, nonanal, and linalool. The remaining odors were presented only during the initial assessment phase to give participants clear points of comparison helping them judge which of the candidate odors felt truly neutral.

The aversive sound was a 500 ms pure tone (1000 Hz) mixed with white noise, and was played at a loudness of 90 dB.

### Apparatus

The odors were presented with the olfactometer (Sniffo, CyNexo S.r.l.). The tubes from the olfactometer were inserted approximately 0.5 cm into both nostrils. The odorants were presented at a combined flow rate of 3 LPM, and were embedded into a continuous stream of odorless air at 1 LPM. A fiber optic cable inserted into one of the olfactory tubes measured the respiration cycle of the participants and determined the point of inhalation with the Spir-O module (Cynexo S.r.l.). The odorant travel time from the jars to the end of the olfactometer tubes was approximately 100 ms.

The aversive sound was presented with an external sound card (Focusrite Saffire Pro 14), and amplifier (Atom Amp+, JDS Labs), via inserted earphones (E-A-RTONE 3A, 3M). Participants used a gamepad (Logitech Gamepad F310) to respond to questions and Visual Analog Scales (VAS). The stimuli, instructions and VAS were presented on a monitor (LG Flatron L227WTP-PF) which was built into the soundproof, electrically shielded cabin.

Participants sat approximately 1 meter away from the monitor, and the light in the cabin was dimmed to 5 lux during the experiment.

### Procedure

First, participants rated the aversive sound and the odors. Based on the rated intensity and pleasantness of the odors, out of the three available odors, the two most neutrally rated odors were chosen as CS. These two odors were then tested on their discriminability in a match-mismatch (discrimination) task. In this task, two odors were presented after each other in four trials. These could either be the same odor or two different odors. Two times the same odor was presented, and two times two different odors were presented with the sequence being different each time. Participants had to rate whether the trial was a match (if the same odor was presented) or mismatch (when two different odors were presented). If participants were not able to correctly identify a match or mismatch on two or more trials, or if they did not identify any of the mismatch trials, then one of the odors was randomly replaced by the remaining neutral odor. All of the participants successfully completed the discrimination task. None of the participants were excluded based on this task.

Conditioning began with an instruction about the contingencies: “In the next part, we will present two odors, and one will be followed by a loud sound. At the end, we will ask you which odor it was.” The first of ten blocks then started. Each block consisted of 15 CS+ and 15 CS-trials (Figure 1). All trials began with a fixation cross that remained on the screen until the VAS. Two seconds after the fixation cross appeared, the odorant was delivered for 500 ms at the participant’s next inhalation. In CS+ trials the 90 dB aversive sound (US) was presented 4 s after odor onset; no sound followed CS-trials. Four seconds after the US offset, participants were prompted to rate the odor’s pleasantness and intensity. After the ratings, an inter-trial interval filled the remaining time so that each trial lasted exactly 25 s.

**Figure 1.**
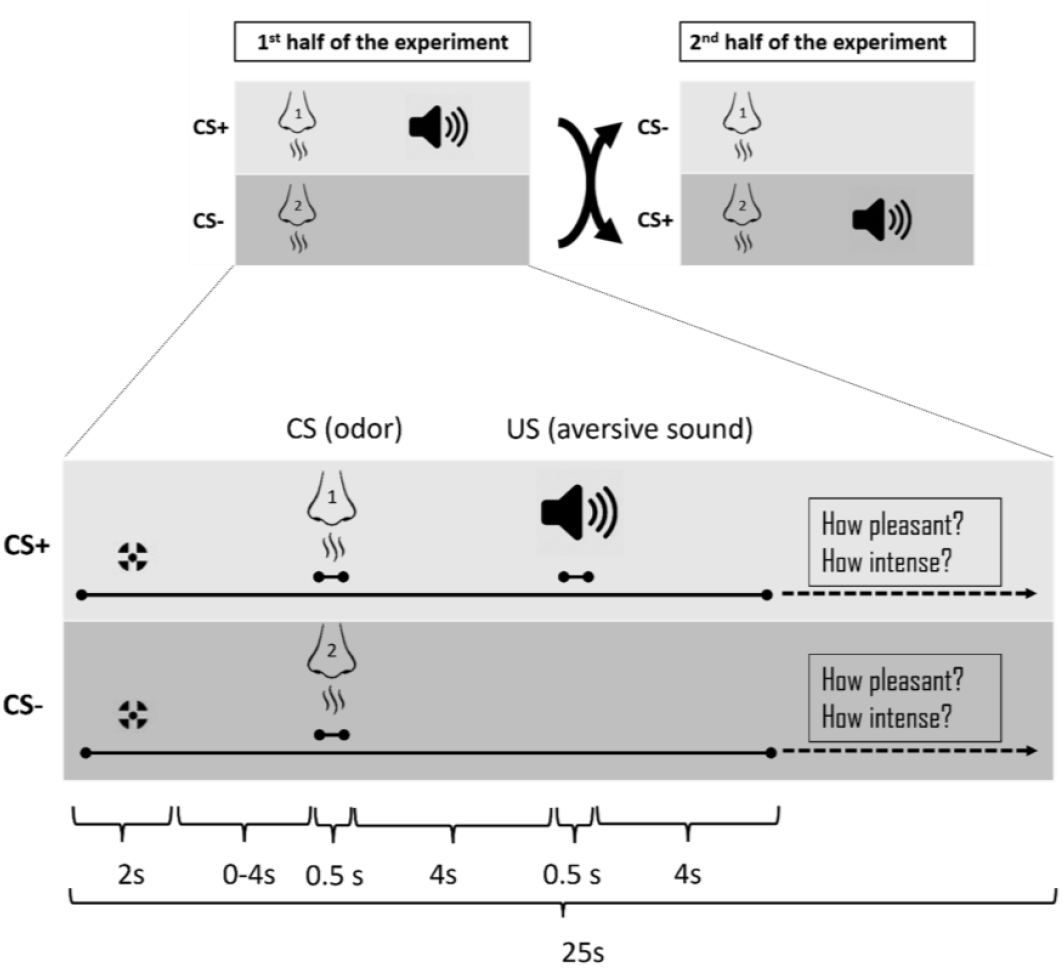
The experimental paradigm. Top panel: the general design of the experiment. In the first half (blocks 1–5) one odor (CS+) was followed by the aversive sound, whereas the other odor (CS−) was not. After five blocks the contingencies were reversed: the initial CS+ was no longer followed by the sound, and the initial CS− was now followed by it. Bottom panel: the trial design. Each trial began with a fixation cross that remained on the screen until the VAS. Two seconds after the fixation cross appeared, the odorant was delivered for 500 ms at the participant’s next inhalation. Four seconds after odor onset, the US was presented in CS+ trials. Four seconds after the US offset, participants rated the odor’s pleasantness and intensity. They then waited during the ITI until the total trial duration reached 25 s.

After five blocks, the CS+ became the CS-, and the CS-became the CS+, i.e., from the sixth block the previously reinforced odor was followed by the aversive sound. The first block and the sixth block (first block after reversal) had a reinforcement rate of 100 %. All other blocks had a reinforcement rate of about 50 %. More specifically, blocks 2, 4, 7 and 9 had eight reinforced CS+ trials each, blocks 3, 5, 8 and 10 had seven reinforced CS+ trials each. The experiment lasted between 4 and 5 hours including the electrode preparation and debriefing.

## Data acquisition and preprocessing

### Behavioral measures

Participants rated the pleasantness (−50 to +50; with anchors “Very unpleasant” and “Very pleasant”) and intensity (0 to 100; with anchors “No sensation” and “Very intense”) of each odor after every trial and rated the US after each block using a visual analogue scale (VAS). We only analyzed the unreinforced CS+ trials for the CS ratings, to prevent the influence of the US presentation on them.

Between task blocks, participants completed a set of questionnaires. This interlude provided a longer break from conditioning, enabled us to monitor changes in state mood, anxiety and sleepiness across the session, and allowed us to characterize the sample on traits relevant to fear learning. German translations of the following instruments were administered via formr (Arslan et al., 2020): State-Trait Anxiety Inventory (Spielberger, 1983), Positive and Negative Affect Schedule (Breyer & Bluemke, 2016), Stanford Sleepiness Scale (Beauducel et al., 2003), Intolerance of Uncertainty Scale (Gerlach et al., 2008), Arnett Inventory of Sensation Seeking (Roth & Mayerhofer, 2003), NEO Five-Factor Inventory (Borkenau & Ostendorf, 2008), and the Behavioral Inhibition System/Behavioral Approach System scales (Strobel et al., 2006). Questionnaire results are presented in Supplementary Material 1.

### Peripheral physiology

Peripheral physiological signals were recorded from AUX channels of the EEG amplifier (actiCHamp, Brain Products GmbH). We used BIP2AUX adapters with a gain of 100, and placed the ground for the physiological channels on the lower right side of the forehead (Figure 2). Data were recorded at a sampling rate of 1000 Hz and filtered offline with a finite impulse response filter using the Hamming window method (mne.filter.filter_data function in MNE-Python; version 1.3.1; (Gramfort et al., 2013)). MNE-Python was used for data preprocessing of all physiological measures and EEG.

**Figure 2.**
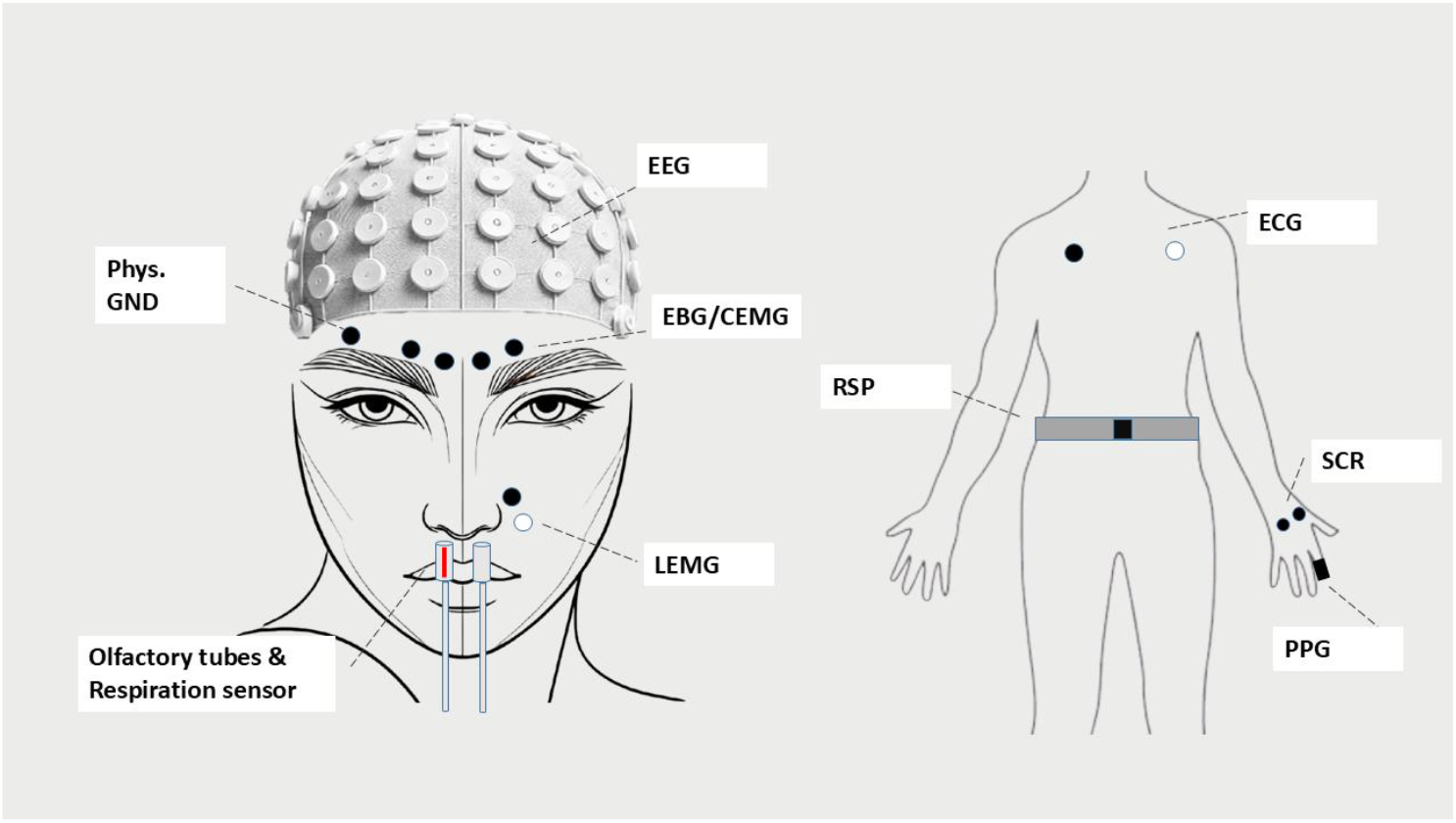
Placement of the sensors. For measures that were recorded with a bipolar montage (LEMG, and ECG), the black dot depicts the positive electrode and white dot the negative electrode. Phys. GND:ground electrode placement for the peripheral physiology measures, EEG: electroencephalography, EBG: electrobulbogram, LEMG: levator labii superioris electromyography, CEMG: corrugator supercilii electromyography, RSP: respiration, ECG: electrocardiography, SCR: skin conductance response, PPG: photoplethysmography.

### Skin conductance response

Due to a technical issue with the skin conductance recording device, only 5 participants remained with sufficient data quality (more than 50 % of trials rejected). We therefore omitted this measure from our analyses.

### Heart rate

ECG was recorded using a BIP2AUX adapter with electrodes on the left and right intersection of the midclavicular and subclavicular line (Figure 2). The signal was filtered with a 0.5 Hz high-pass filter, a 30 Hz low-pass filter, and a 50 Hz notch filter to improve R-peak detection. The data were epoched into initial [-4000 – 11,000 ms] bins, where 0 represented the CS onset. Automatic R-peak detection with manual correction was performed with HRVTool (Vollmer, 2019). R-peaks were then transformed to RR intervals, and then converted to instantaneous beats per minute (BPM). After that, the epochs for analysis were cropped into [-1500 – 4000 ms] bins. The reason for initially having larger bins was to account for long RR intervals at the epochs’ edges, and to inspect responses to the US. BPM values above 220 or below 50 were removed from the data. Then, the epoch was divided into 500 ms bins (i.e. 12 bins with an epoch length of 5.5 s), and for each bin we checked whether one or multiple BPM values were present. If only less than half of the bins contained a BPM value, the trial was rejected. Otherwise, the signal was interpolated to create an epoch containing one BPM value per bin, and offering a temporal development of the heart rate. The resulting epochs were then baseline-corrected per participant by calculating the average baseline signal for each CS condition, and subtracting it from the signal. We then analyzed the average heart rate change over the first acceleration after CS onset [500 – 2000 ms], the second deceleration after CS onset [2000 – 4000 ms], and the acceleration in the time window [5000 – 8000 ms] after CS onset. The latter analysis was conducted to examine responses to the absence of the US in unreinforced CS+ trials (Sperl et al., 2021).

### Pulse wave amplitude

Photoplethysmography (PPG) was measured using a PPG module (FpSens A1; MKS) on the left index finger (Figure 2). The data were filtered with a 0.5 Hz high-pass filter and a 15 Hz low-pass filter. The data were epoched in the same way as the heart rate (see above), where originally larger epochs were created than the ones used for the analysis. The systolic peak and the preceding base were detected with HRVTool (Vollmer, 2019). The pulse wave amplitude was then calculated by subtracting the base from the peak amplitude. A peripheral vasoconstriction response associated with attenuation of pulse wave amplitude is associated with increased sympathetic activity (Babchenko et al., 2001). Values above or below 3 *SD* of the mean were removed from the data. Then, similar to the heart rate preprocessing, we created 500 ms bins, and interpolated the pulse wave amplitude values in each epoch, rejecting trials if less than half of the bins contained pulse wave amplitude values. Participants with more than half of the trials rejected were excluded from analyses. The pulse wave amplitude was then corrected by calculating the percentage increase to the baseline. The pulse wave amplitude was analyzed over an initial period of pulse wave amplitude decrease [500 – 2000 ms], and a subsequent increase [2000 – 4000 ms] after CS onset. Similar to the heart rate analysis, the time window [4000 – 8000 ms] after CS onset was also analyzed to examine the effect of absence of the US in unreinforced trials.

### Respiration

Respiration was measured using a respiration belt (Brain Products GmbH). The respiration belt was placed around the waist between the lowest rib and the navel (Figure 2). A 5 Hz low-pass filter was applied to the raw signal, and it was manually inspected for artifacts after epoching the data into initial [-2000 – 6000 ms] bins, where 0 represents the CS onset. Each epoch was baseline corrected, by subtracting the average of the baseline. The respiration amplitude was calculated by averaging over the inhalation peak after CS onset, which was in the time window of [1000 – 2000 ms] after CS onset. The respiration rate was calculated using Neurokit2 (version 0.2.10), where the inter-breath-intervals were calculated for initial [-10,000 – 20,000 ms] epochs in order to also correctly analyze the respiration rate at the edges of the epochs. The epoch was then cropped into [-2000 – 6000 ms] bins. The resulting data was baseline corrected by subtracting the average baseline, and then analyzed by averaging over the time window after CS onset [0 – 4000 ms].

### Levator EMG

The levator labii superioris EMG was measured using a BIP2AUX adapter, placing one electrode on the middle of the levator labii superioris alaeque nasi on the left side of the nose, and the other electrode below it (Figure 2). The raw signal was first filtered with a 20 Hz high-pass filter, 499 Hz low-pass filter, and a 50 Hz notch-filter. The filtered signal was then epoched into initial [-1000 – 5000 ms] bins, and all trials with a peak-to-peak signal amplitude higher than 30 mV were rejected automatically. This liberal threshold was based on the suggestion by (Raez et al., 2006) that an EMG signal has a range of 0 – 10 mV (+5 or -5). The remaining trials were then rectified, and a 50 Hz low-pass filter was applied. Following the recommendations of (Rutkowska et al., 2023, p. 202), the muscle activity of the levator muscle was then baseline-corrected by dividing by the average baseline amplitude. The response was measured as the mean amplitude in the time window [0 – 4000 ms] after CS onset.

### Corrugator EMG

The muscle activity at the left corrugator supercilii was extracted from the EEG electrodes that were placed there initially to measure the electrobulbogram (EBG; (Iravani et al., 2020); Figure 2). The electrode closest to the midline of the face above the left eyebrow was used for the signal, and the electrode further away from the midline above the left eyebrow was used as reference. After recording, the signal was re-referenced to this reference. The raw signal was then pre-processed in the same way as the levator EMG. Similar to the levator EMG, the signal of the corrugator EMG was baseline-corrected by dividing by the average baseline amplitude. Responses are the mean muscle amplitude in the time window [0 – 4000 ms] after the CS onset.

### EEG

We recorded EEG with the actiCHamp 64 channel system (Brain Products GmbH). Electrodes were placed according to the 10-20 system with Cz channel as the online reference and Fpz as the ground electrode. We placed four electrodes (PO3, PO4, PO7, and PO8) above the eyebrows for an EBG (Iravani et al., 2020). Impedances were maintained below 25 kOhm. The sampling rate was 1000 Hz.

Each recording was filtered offline by applying 0.1 Hz high-pass and 45 Hz low-pass filters. Data were re-referenced to average reference and epoched into [-2000 – 5000 ms] bins with 0 representing the CS onset. Epochs containing high-amplitude artifacts that would distort the Independent Component Analysis (ICA) were visually identified and discarded. Noisy channels were excluded to later be interpolated, and then, an ICA was performed using the Picard algorithm. As a preliminary step for improving ICA decomposition, a 1 Hz high-pass filter was applied. The results of the ICA on the 1 Hz filtered data were then imported to the data filtered with the 0.1 Hz high-pass filter. Components clearly related to blinks, eye movements and high amplitude muscle activity were removed. Then, the excluded channels were interpolated (interpolate_bads function in MNE-Python), and reintegrated in the dataset.

### Event-related potentials

The late positive potential (LPP) can reflect the valence of an odor (Lundström et al., 2006). (Lundström et al., 2006) located this component, to which they refer as P3, in the time window around 825 to 927 ms after odor onset. As odor latencies may vary, we report the difference in average amplitude in the time window [750 – 1000 ms] at the Pz electrode. In addition we analyzed the average amplitude in the time window [3800 – 4000 ms] after CS onset at the Cz electrode, as in other studies the interval of 100 to 200 ms before US onset is chosen to measure the SPN (Böcker et al., 2001; Tanovic & Joormann, 2019). For these analyses the EEG signal was re-referenced to the average of both mastoid electrodes (TP9 and TP10).

### Time-frequency analysis

To assess differences in alpha suppression between conditions, we conducted a time-frequency analysis using Morlet wavelets (using the mne.time_frequency.tfr_morlet function, 1 to 45 Hz, equally spaced, and number of cycles being half of each frequency). The EEG was re-referenced to the average of all electrodes. Permutation-based clustering (using the permutation_cluster_1samp_test function) in the alpha band (8 – 12 Hz) from 1.5 to 4 s after CS onset was used to test which electrodes exhibited a significant signal change compared to baseline activity. This method revealed two clusters with significantly stronger alpha suppression: one over left centro-parietal electrodes (C3, CP1, C5, CP3) and another over right centro-parietal electrodes (CP6, CP2, C4, T8, CP4, C6). We used these electrodes for further statistical analyses.

### Temporal decoding

Because olfactory inputs originate in the olfactory bulb and project diffusely to the piriform cortex rather than to a single, well-defined cortical source (Sosulski et al., 2011), the scalp distribution of CS+ minus CS-ERPs can differ considerably from one participant to the next. Such spatial variability makes it suboptimal to rely on fixed electrode locations such as Fz, Cz, or Pz. To accommodate these differences, we applied within-participant temporal decoding to two feature sets: the full time-domain ERP and alpha-band power extracted from the TFR.

EEG data were downsampled to 100 Hz and epoched from 0 to 4000 ms relative to CS onset. For each participant we trained a support vector machine classifier (sklearn.svm.SVC) to distinguish CS+ from CS-trials. Features were z-scored with sklearn.preprocessing.StandardScaler, and decoding was performed with MNE-Python’s SlidingEstimator, which fits a separate classifier at every time point and evaluates performance using the area under the ROC curve (AUC). Ten-fold cross-validation was conducted within participants, and AUCs were averaged across folds to generate an accuracy curve for each individual. Group-level decoding was obtained by averaging these curves across participants at every time point.

## Statistical analysis

We used the Pingouin package (version 0.5.3) to conduct ANOVAs. In all measures, except the CS and US ratings, we analyzed the factor CS with a two-way RM ANOVA (Condition x Reversal: with two levels of the factor Condition: CS+, CS-, and two levels of the factor Reversal: before reversal, after reversal). For the respiration, EMG and EEG analyses, we averaged the unreinforced and reinforced CS+ trials. For heart rate and pulse wave amplitude, we analyzed reinforced and unreinforced CS+ trials separately, thus the factor Condition consisted of three levels: CS+reinforced, versus CS+unreinforced, versus CS-.

For analysis of the CS ratings, we performed a similar analysis, however did not analyze the reinforced trials, as these could influence the CS ratings which were given after the US onset. Therefore, the CS ratings were analyzed with a two-way RM ANOVA (Condition x Reversal: with two levels of the factor Condition: CS+unreinforced, and CS-, and two levels of the factor Reversal: before reversal, versus after reversal).

For analysis of the US ratings, we used a one-way RM ANOVA (factor Block with 11 levels of block) to analyze the habituation of the US over the course of the experiment.

Considering our reversal paradigm, and our analysis approach where only the condition (CS+ odor, versus CS-odor), and not the odorant (odor 1, versus odor 2) are included as factors, the RM ANOVA should be interpreted as follows. A significant main effect of Condition (CS+ vs. CS-) indicates successful conditioning across the experiment, including both initial acquisition and relearning after reversal. A Condition × Reversal interaction would suggest that the CS difference changed after the reversal. If a significant difference between CS+ and CS-is present only before the reversal - or if the difference persists in the direction of the original odor after the reversal (i.e., the initial CS+ still elicits stronger responses despite being the new CS-) - this would indicate a failure to relearn. In contrast, a main effect of Reversal without an interaction is less informative for interpreting learning and likely reflects general temporal effects such as habituation across the session.

The assumption of sphericity was violated in some analyses, and in that case greenhouse-geisser corrected *p*-values are reported. The *p*-values reported in post-hoc comparison *t*-tests have been corrected with the Benjamin-Hochberg FDR. Also, due to the exploratory nature of our study, the significance threshold for ANOVAs was set at *p* < .01.

The results of the temporal decoding of the ERP were analyzed using permutation-based clustering (mne.stats.spatio_temporal_cluster_1samp_test in MNE-Python; 1024 permutations; *t*-threshold was determined with the scipy.stats.rv_continuous.ppf function) to identify clusters in time where accuracy scores were significantly (*p* < .05) above chance level (50 %).

## Results

### Behavioral

#### CS ratings

Analysis of the pleasantness ratings revealed no significant difference between the unreinforced CS+ and CS-, *F*(1, 29) = 1.11, *p* = .30, η_p_^2^ = 0.04, no significant main effect of reversal, *F*(1, 29) = 2.13, *p* = .16, η_p_^2^ = 0.07, and no interaction effect was found either, *F*(1, 29) = 0.04, *p* = .84, η _p_^2^ < 0.01. Figure 3 shows that the mean ratings of both odors were neutral before and remained neutral after reversal, but that the second odor (i.e., the one that was CS-before and CS+ after reversal) had a broader distribution of ratings than the first odor.

**Figure 3.**
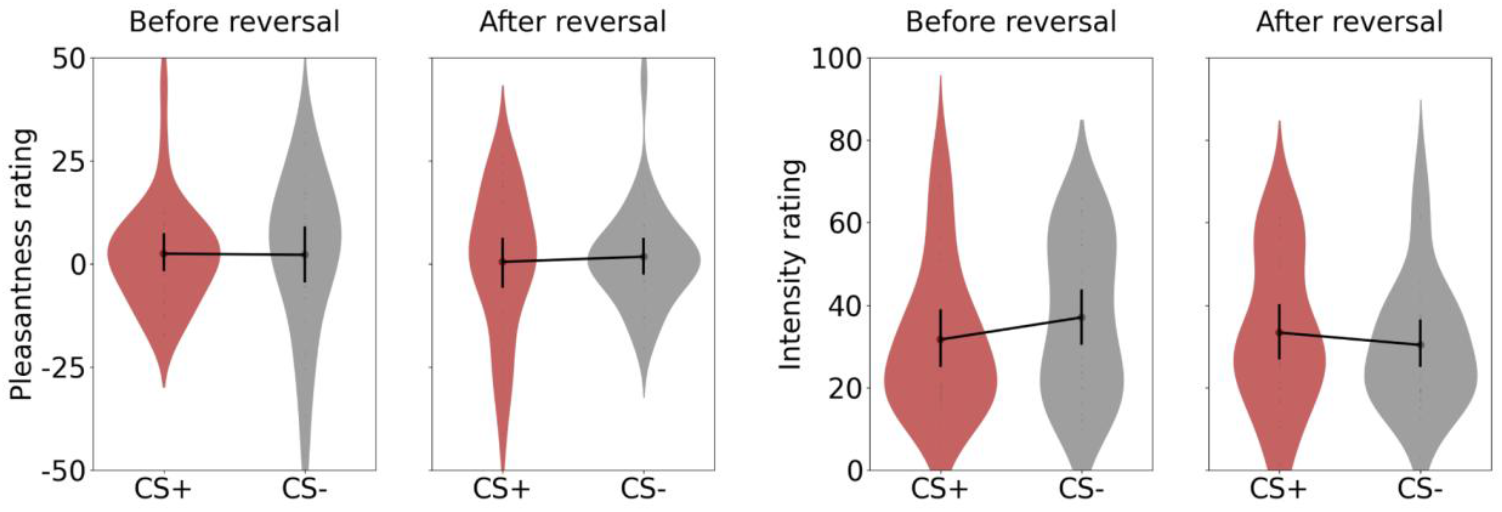
Pleasantness and intensity ratings of the CS before and after reversal. Error bars represent 95% CI.

Similarly, analysis of the intensity ratings revealed no significant effects of condition, *F*(1, 29) = 2.03, *p* = .16, η_p_^2^ = 0.07, of reversal, *F*(1, 29) = 4.65, *p* = .04, η_p_^2^ = 0.14, or interaction effect, *F*(1, 29) = 1.76, *p* = .19, η_p_^2^ = 0.06 (Figure 3).

### US ratings

On the scale of -50 (very unpleasant) to +50 (very pleasant), the US (aversive sound) was rated as unpleasant on average (*M* = -26,29, *SD* = 14.62). Similarly, the US was rated as rather intense on the scale of 0 – 100 (*M* = 67.13, *SD* = 19.13). US ratings remained stable across blocks, as shown by the non-significant effects of Block on Pleasantness, *F*(10, 290) = 1.25, *p* = .29, η_p_^2^ = 0.04, and Intensity, *F*(10, 290) = 1.76, *p* =.12, η_p_^2^ = 0.06.

### Peripheral physiology

#### Heart rate

Analysis of the first acceleration in the heart rate (0.5 – 2 s time window; see Figure 4) showed no significant main effect of CS, *F*(2, 46) = 0.17, *p* = .79, η_p_^2^ = 0.01, nor was there a main effect of reversal, *F*(1, 23) = 0.25, *p* = .62, η_p_^2^ = 0.01, or interaction effect of CS and reversal, *F*(2, 46) = 1.58, p = .22, η_p_^2^ = 0.06. The second deceleration (2 – 4 s) also showed no main effect of CS, *F*(2, 46) = 2.15, *p* = .14, η_p_^2^ = 0.09, main effect of reversal, *F*(1, 23) = 0.05, *p* = .83, η_p_^2^ < 0.01, or interaction effect, *F*(2, 46) = 0.02, *p* = .97, η_p_^2^ < 0.01.

**Figure 4.**
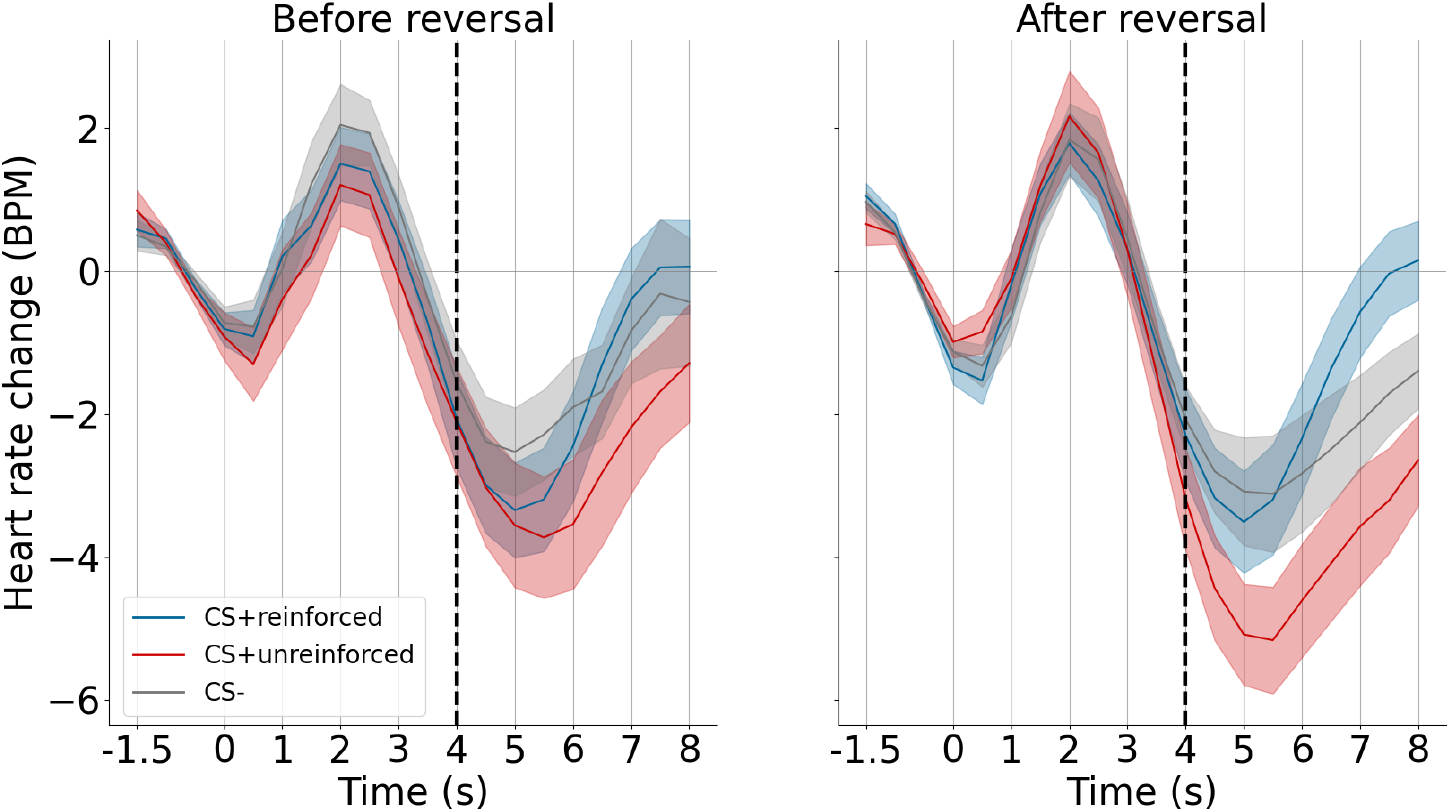
Heart rate response to the CS before and after reversal. The dashed line indicates the US onset in CS+reinforced trials. Shaded areas represent the standard error of the mean (SEM).

The post-US acceleration (5 – 8 s) revealed a significant main effect of condition, *F*(2, 46) = 8.57, *p* = .002, η_p_^2^ = 0.27, but no main effect of reversal, *F*(1, 23) = 1.45, *p* = .24, η_p_^2^ = 0.06, and no interaction effect, *F*(2, 46) = 1.76, *p* = 0.19, η_p_^2^ = 0.07. Post-hoc comparisons showed a significant difference between reinforced and unreinforced CS+ trials, *t*(23) = 3.43, *p* = .007, *d* = 0.57, however, no difference between reinforced CS+ and CS-trials (*p* = .04, *d* = 0.24), and no difference between unreinforced CS+ and CS-trials (*p* = .02, *d* = -0.36).

### Pulse wave amplitude

The initial decrease in pulse wave amplitude was not different between conditions, *F*(2, 32) = 0.92, *p* = .40, η_p_^2^ = 0.05, and no main effect of reversal, *F*(1, 16) = 2.39, *p* = .14, η_p_^2^ = 0.13, or interaction effect was significant, *F*(2, 32) = 0.04, *p* = .89, η_p_^2^ < 0.01 (Figure 5). Similarly, no main effect of condition was found for the subsequent increase of the pulse wave amplitude, *F*(2, 32) = 0.97, *p* = .39, η_p_^2^ = 0.06, and no main effect of reversal, *F*(1, 16) = 4.05, *p* = .06, η_p_^2^ = 0.20, or interaction effect, *F*(2, 32) = 1.80, *p* = .19, η_p_^2^ = 0.10.

**Figure 5.**
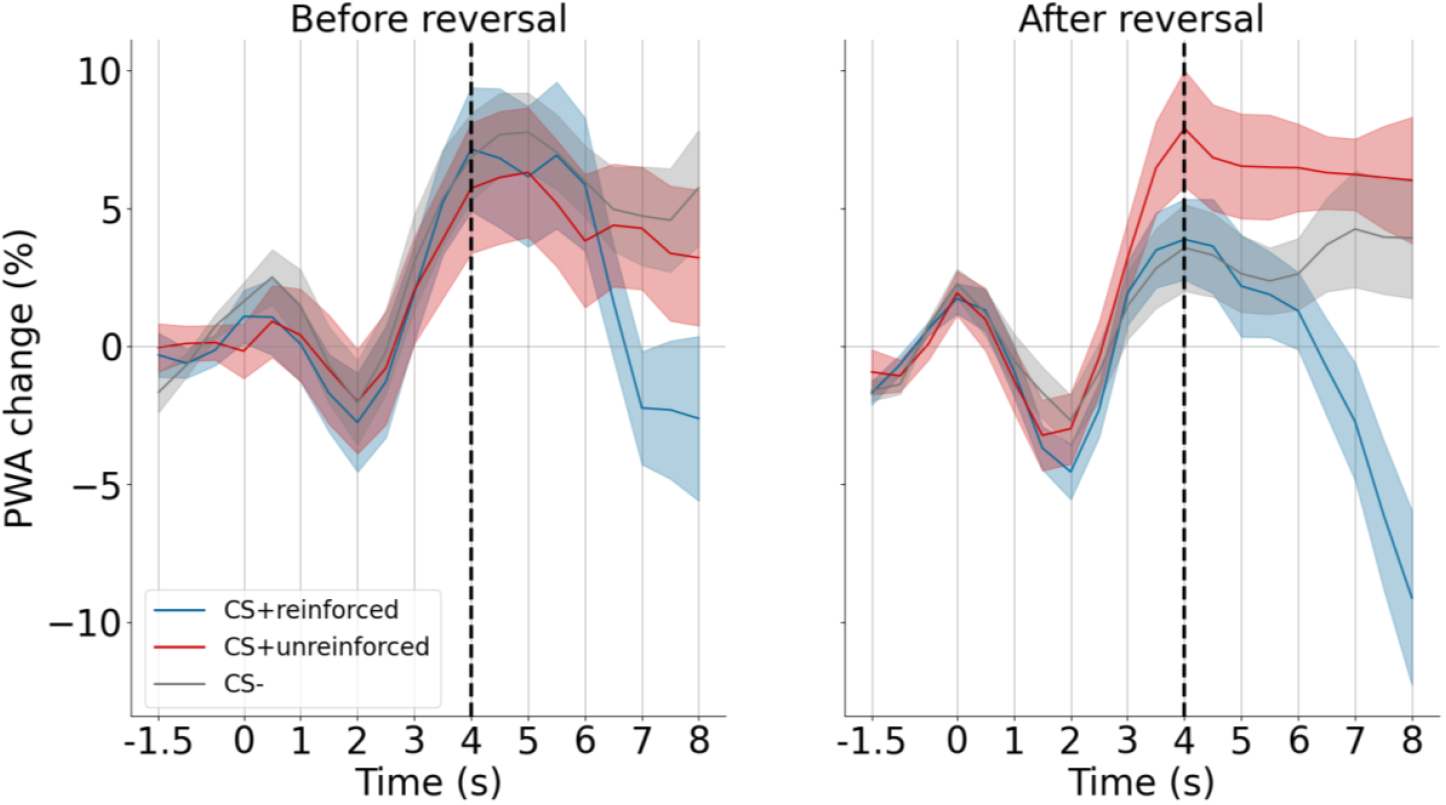
Percentual change of the pulse wave amplitude after the CS onset compared to baseline before and after reversal. The dashed line indicates the US onset in CS+reinforced trials. Shaded areas represent the SEM.

In the post-US time window a significant main effect was found for condition, *F*(2, 32) = 7.25, *p* = .005, η_p_^2^ = 0.31, however no main effect of reversal, *F*(1, 16) = 1.07, *p* = .32, η_p_^2^ = 0.06, or interaction effect, *F*(2, 32) = 3.04, *p* = .06, η_p_^2^ = 0.16. Post-hoc comparisons showed, however, that there were no significant differences between the reinforced CS+ and unreinforced CS+ (*p* = .02, *d* = -0.77) or CS-(*p* = .02, *d* = -0.68). Neither was there a significant difference between the unreinforced CS+ and the CS-(*p* = .44, *d* = 0.14).

#### Respiration

Respiration amplitude between 1 and 2 s after CS onset showed no significant main effect of CS, *F*(1, 29) = 0.05, *p* = .82, η_p_^2^ < 0.01, nor an effect of reversal, *F*(1, 29) = 1.37, *p* = .25 η_p_^2^ = 0.05. No interaction effect was revealed either, *F*(1, 29) = 1.01, *p* = .32, η_p_^2^ = 0.03.

The respiration rate showed no main effect of CS either, *F*(1, 28) = 0.98, *p* = .33, η _p_^2^ = 0.03, no main effect of reversal, *F*(1, 28) = 1.95, *p* = .17, η _p_^2^ = 0.06, and no interaction effect of CS and reversal, *F*(1, 28) = 0.20, *p* = .66, η _p_^2^ = 0.01.

#### Levator EMG

The average levator EMG amplitude showed no significant differences between the different CS conditions, *F*(1, 29) = 0.03, *p* = .87, η_p_^2^ < 0.01. Nor was there an effect of reversal, *F*(1, 29) = 4.22, *p* = .05, η_p_^2^ = 0.13 or an interaction effect of condition and reversal, *F*(1, 29) = 2.70, *p* = .11, η_p_^2^ = 0.09.

#### Corrugator EMG

The corrugator EMG showed no differences in response to the CS, *F*(1, 29) = 1.18, *p* = .29, η_p_^2^ = 0.04. Likewise, no main effect of reversal existed, *F*(1, 29) = 0.03, *p* = .87, η_p_^2^ < 0.01, and the interaction effect was not significant, *F*(1, 29) = 1.30, *p* = .26, η_p_^2^ = 0.04.

## EEG

### ERP

Analysis of the LPP did not show a significant main effect of condition, *F*(1, 27) = 0.15, *p* = .70, η_p_^2^ < 0.01, no significant main effect of reversal, *F*(1, 27) = 0.65, *p* = .43, η_p_^2^ = 0.02, and no interaction effect, *F*(1, 27) < 0.01, *p* = .99, η_p_^2^ < 0.01 (Figure 6).

**Figure 6.**
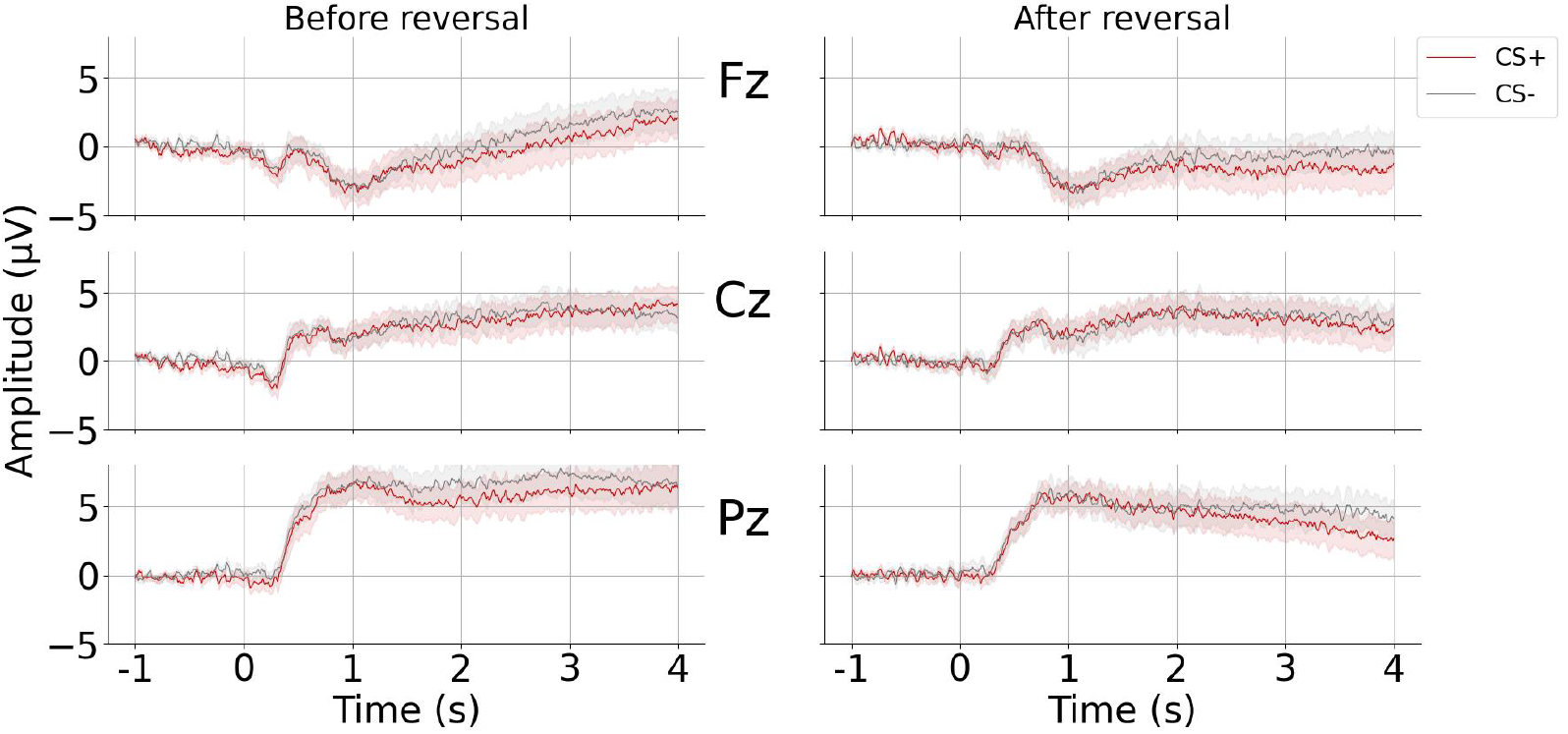
ERPs to the CS before and after reversal at the electrodes Fz, Cz, and Pz. The CS+ represents the average of the CS+reinforced and CS+unreinforced trials. Shaded areas represent the SEM.

Analyzing the SPN showed that the main effect of condition was not significant, *F*(1, 27) < 0.01, *p* = .98, η_p_^2^ < 0.01, neither was the main effect of reversal significant, *F*(1, 27) = 2.61, *p* = .12, η_p_^2^ = 0.09, or the interaction effect of condition and reversal, *F*(1, 27) = 0.68, *p* = .42, η_p_^2^ = 0.02.

#### Time-frequency analysis

Analysis of the alpha suppression at centro-parietal electrodes over the left hemisphere showed no significant main effect of CS, *F*(1, 27) = 3.50, *p* = .07, η_p_^2^ = 0.11, nor was a significant main effect of reversal present, *F*(1, 27) = 6.61, *p* = .016, η_p_^2^ = 0.20, or an interaction effect, *F*(1, 27) = 0.04, *p* = .84, η_p_^2^ < 0.01 (Figure 7). Similarly, centro-parietal electrodes over the right hemisphere showed no differences in alpha suppression between CS conditions, *F*(1, 27) = 2.17, *p* = .15, η_p_^2^ = 0.07, and no significant interaction effect of CS and reversal, *F*(1, 27) < 0.01, *p* = .99, η_p_^2^ < 0.01. However, there was a significant main effect of reversal, *F*(1, 27) = 10.02, *p* = .004, η_p_^2^ = 0.27. Post-hoc comparisons, however showed a similar difference between before and after reversal for CS+ trials, *t*(27) = 3.07, *p* = .005, *d* = 0.18, as for CS-trials, *t*(27) = 3.09, *p* = .005, *d* = 0.19.

**Figure 7.**
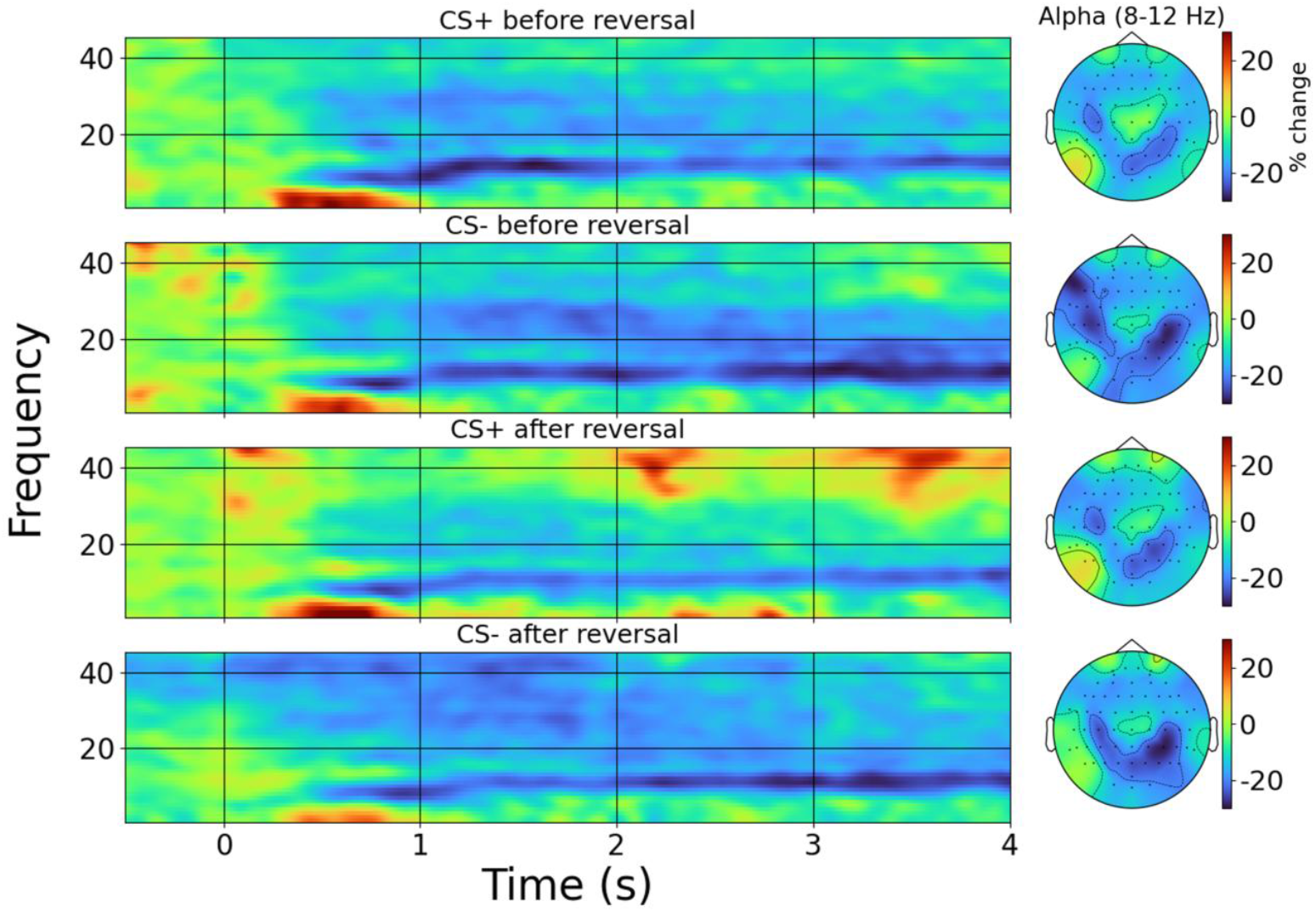
Time-frequency representations (TFR) and topographic plots before and after reversal at the centro-parietal electrodes combined for the clusters of the left and right hemisphere. The topographic plots represent the average in the time window [1500 – 4000 ms] and alpha frequency band (8 – 12 Hz). Color bars indicate the percentage signal change to the baseline in both the TFRs and topographic plots.

#### Temporal decoding

Analysis of the decoding classifier on ERP data showed two clusters of time points after CS onset where the classifier performed better than chance level. One cluster (*p* = .015) was identified for the time points [2240 – 2870 ms], where the average accuracy was 53.02 % (*SD* = 0.59). Another cluster (*p* = .005) was identified for the time points [2910 – 4000 ms], where the average accuracy was 53.45 % (*SD* = 0.46).

Analysis of the decoding classifier on TFR data averaged in the alpha frequency band showed three clusters of time points. One cluster (*p* = .03) was found for the time points [2740 – 3040 ms], with an average decoding accuracy of 52.90 % (*SD* = 0.39). Another cluster (*p* = .003) was found for the time points [3200 – 3620 ms], where the average accuracy was 53.33 % (*SD* = 0.73), and one cluster (*p* = .02) in the time window [3720 – 4000 ms], where the average accuracy was 53.12 % (*SD* = 0.53).

## Discussion

This study investigated the behavioral, physiological, and neural correlates of olfactory reversal conditioning using a broad range of measures. Despite extensive data collection, we observed no significant differences between CS+ and CS-in behavioural ratings, autonomic responses, facial electromyography, or univariate EEG components.

Multivariate analyses yielded a limited exception. Support vector machine classifiers trained on single-trial time-domain EEG and on alpha-band power distinguished CS+ from CS− at above-chance levels during late post-stimulus intervals (> 2.2 s), with mean accuracies of approximately 53 %. The reason why we found an effect here, but not in the grand-averaged ERPs or with TFR, could be the large interindividual differences in the spatial distribution of EEG. The limited sensitivity of our physiological measures aligns with other studies that did not find strong differences in odor-shock pairings (Cavazzana et al., 2018; Li et al., 2008) or odor-CO_2_ pairings (Bensafi et al., 2007; Moessnang et al., 2013).

One possible explanation for the lack of successful conditioning is that the CS was not sufficiently discriminable. However, we took steps to address this by including a discrimination task before conditioning and replacing the odor that participants could not adequately distinguish. In addition, other olfactory conditioning studies have shown that participants often develop enhanced sensitivity (Åhs et al., 2013; Parma et al., 2015) and improved discrimination between odors (Li et al., 2008; Pool et al., 2014) after conditioning. These findings make it unlikely that poor CS discriminability alone explains the lack of conditioning in our study.

Another potential factor is the duration of the experiment, which could lead to fatigue or habituation and weaken responses over time. Yet, a block-wise analysis revealed no differences between CS+ and CS-in the earliest blocks, and the US remained both unpleasant and intense throughout the experiment. These observations suggest that fatigue or habituation cannot fully account for the unsuccessful conditioning.

A more plausible explanation may lie in the temporal structure of the paradigm. Our use of a 3.5 second trace interval between odor offset and US presentation may have disrupted the formation of strong associations. Although some studies have reported successful trace conditioning with intervals up to 2.6 seconds (Howard et al., 2016; Moessnang et al., 2013; You et al., 2022), others using similar or even shorter intervals have failed to detect robust effects (Bensafi et al., 2007; Cavazzana et al., 2018; Li et al., 2008). This inconsistency suggests that olfactory conditioning may be more sensitive to timing and procedural details than visual or auditory modalities.

These results also raise broader questions about the generalisability of classical conditioning paradigms across sensory domains. The olfactory system’s unique anatomy - characterised by minimal thalamic relay and direct access to limbic structures - may favour innate affective responses over rapid associative plasticity. Additionally, pairing olfactory CS with auditory US may result in weaker cross-modal associations compared to within-modality conditioning.

Overall, our findings suggest that olfactory conditioning may require more precise methodological calibration than conditioning in other sensory modalities. Systematic comparisons across experimental designs are needed to identify the conditions under which robust learning occurs. Establishing these boundary conditions will be critical for advancing theoretical models of associative learning and for developing translational paradigms relevant to clinical interventions targeting maladaptive affective associations.

## Acknowledgments

This study was supported by the Else-Kröne-Fresenius Stiftung. We would further like to thank Simone Liebscher at the Schenke-Layland lab in Tübingen for her help in creating the olfactory stimuli.

## Data and code availability statement

All raw data and code of the experiment can be retrieved at https://gin.g-node.org/nickmenger/Menger_et_al_2025_Olfactory_reversal_conditioning.git. The behavioral data can be retrieved at https://osf.io/dtck2/

## CRediT statement

**NS Menger**: Conceptualization, Methodology, Software, Validation, Formal analysis, Investigation, Resources, Data curation, Writing - original draft, Writing - review & editing, Visualization

**YG Pavlov**: Conceptualization, Methodology, Software, Validation, Resources, Writing - review & editing, Supervision, Project administration

**B Kotchoubey**: Conceptualization, Methodology, Writing - review & editing, Supervision, Project administration, Funding acquisition

## Supplementary material

### SM 1. Questionnaire results

**Table 1.** Mean (*SD*) personality trait scores.

| Sample size | STAI trait | IUS factor 1 | IUS factor 2 | AISS novelty | AISS intensity | NEO (N) | NEO (E) | NEO (O) | NEO (C) | NEO (A) | BIS/BAS factor 1 | BIS/BAS factor 2 | BIS/BAS factor 3 | BIS/BAS factor 4 |
| --- | --- | --- | --- | --- | --- | --- | --- | --- | --- | --- | --- | --- | --- | --- |
| $n = 29$ | 36.5 (8.3) | 60.2 (17.9) | 53.5 (16.8) | 2.7 (0.4) | 2.3 (0.4) | 21.5 (6.2) | 25.6 (6.0) | 18.3 (3.5) | 29.9 (5.0) | 16.8 (2.7) | 11.1 (2.8) | 11.1 (2.9) | 15.6 (3.7) | 19.4 (5.2) |

*Note:* STAI: State-Trait Anxiety Inventory, IUS: Intolerance of Uncertainty Scale, AISS: Arnett Inventory of Sensation Seeking, NEO: NEO Five-Factor Inventory, BIS/BAS: Behavioral Inhibition System and Behavioral Approach System.

Before, during, and after the experiment, participants rated their anxiety level (Figure 1). The STAI state anxiety level did not significantly change over the duration of the experiment, *F*(2, 54) = 4.73, *p* = .013, η_p_^2^ = 0.15.

**Figure 1.**
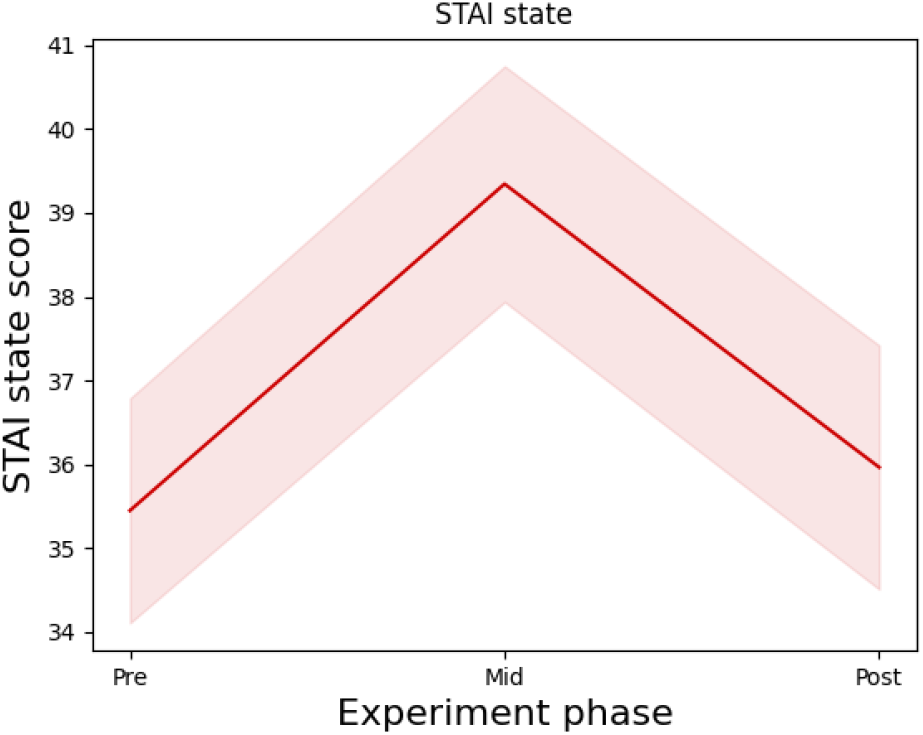
STAI state scores over time. Shaded areas represent the SEM.

The sleepiness of participants was measured after every other block, and showed significant changes over time (Figure 2), *F*(5, 135) = 11.87, *p* < .001, η _p_^2^ = 0.31. Post-hoc comparisons revealed a significant increase between the pre-experiment sleepiness score, and all other scores (all *p* < .01), and no further significant differences.

**Figure 2.**
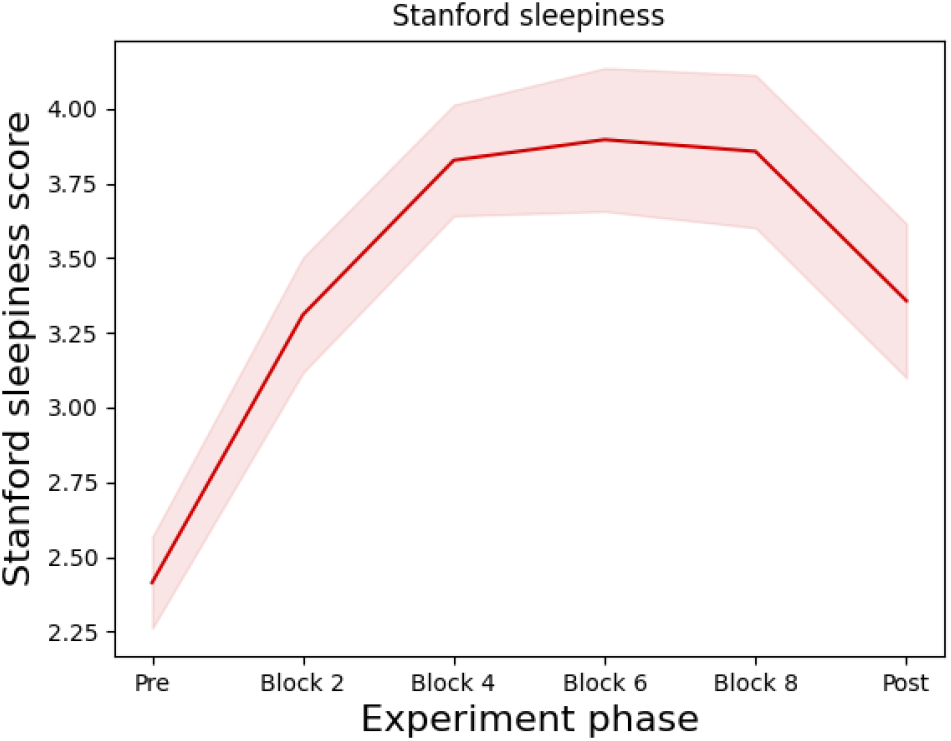
Stanford sleepiness scores over time. Shaded areas represent the SEM.

The positive and negative affect were measured before, during and after the experiment (Figure 3). The positive effect differed significantly, *F*(2, 54) = 13.47, *p* < .001, η_p_^2^ = 0.33, which showed a decrease during the experiment compared to before (*p* < .001), but did not differ during or after the experiment (*p* >= .019). The negative affect showed no significant difference, *F*(2, 54) = 2.21, *p* = .12,η_p_^2^ = 0.08.

**Figure 3.**
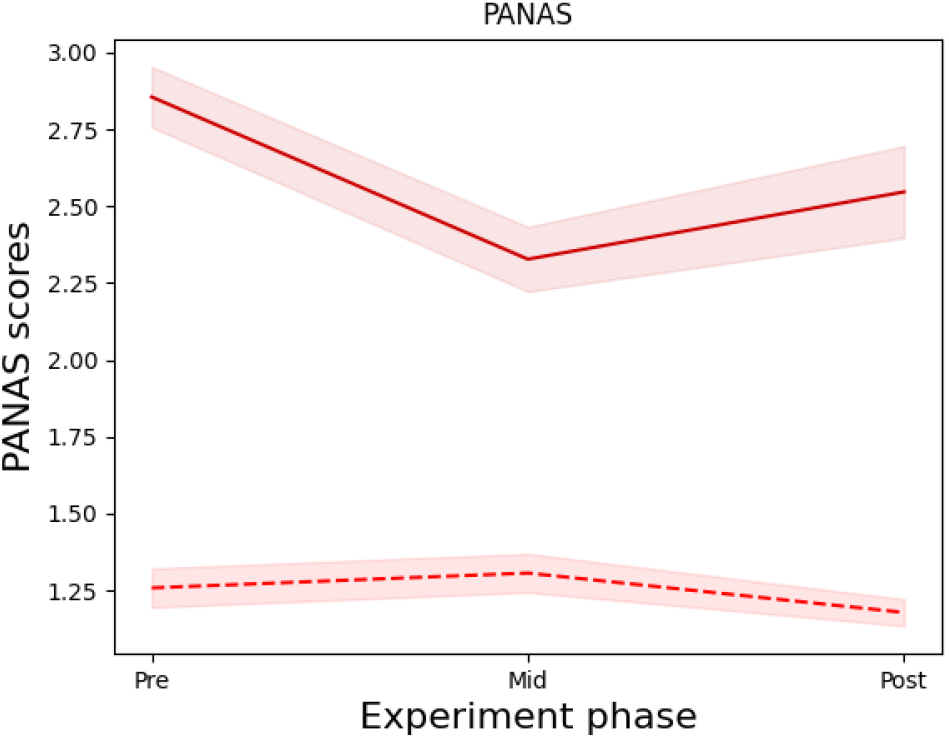
Positive affect (solid) and negative affect (dashed) scores over time. Shaded areas represent the SEM.

